# About bacteriophage tail terminator and tail completion proteins: structure of the proximal extremity of siphophage T5 tail

**DOI:** 10.1101/2024.10.16.618446

**Authors:** Romain Linares, Cécile Breyton

## Abstract

Bacteriophages are viruses infecting bacteria. The vast majority of them bear a tail, allowing host recognition, cell wall perforation and DNA injection into the host cytoplasm. Using electron cryo-microscopy (cryo-EM) and single particle analysis, we determined the organisation of the tail proximal extremity of siphophage T5 that possess a long flexible tail, and solved the structure of its tail terminator protein (TrP) p142 (TrP_142_). It allowed to confirm the common evolutionary origin between T5 TrP_p142_ and other known or putative TrPs from siphophages, myophages and bacterial tail-like machines, despite very poor sequence conservation. By also determining the structure of T5 tail proximal extremity after interaction with T5 bacterial receptor FhuA, we showed that no conformational changes occur in TrP_p142_ and confirmed that the infection signal transduction is not carried by the tube itself. We also investigated the location of T5 tail completion protein (TCP) p143 (TCP_p143_) and showed, thanks to a combination of cryo-EM and structure prediction using Alphafold2, that it is not located at the capsid-to-tail interface as suggested by its position in the genome, but instead, very unexpectedly, on the side of T5 tail tip, and that it appears to be monomeric. Based on structure comparison with other putative TCPs predicted structures, this feature could not be shared by other TCPs. The stoichiometry of the Tape Measure Protein is also discussed.

**Importance:** Bacteriophages, viruses infecting bacteria, are the most abundant living entities on Earth. They are present in all ecosystems where bacteria develop and are instrumental in the regulation, diversity, evolution and pathogeny of microbial populations. Moreover, with the increasing number of pathogenic strains resistant to antibiotics, virulent phages are considered as a serious alternative or complement to classical treatments. 96% of all phages present a tail that allows host recognition and safe channelling of the DNA to the host cytoplasm. We present the atomic model of the proximal extremity of siphophage T5 tail, confirming structural similarities with other phages. This structure, combined to results previously published further explored, also allowed a review and a discussion on the role and localisation of a mysterious tail protein, the Tail Completion Protein, which is known to be present in the phage tails, but that was never identified in a phage structure.

## Introduction

Bacteriophages – phages for short – are viruses infecting bacteria. They represent the most abundant biological entity on our planet. A large majority of them are composed of a capsid filled with double-stranded DNA and of a tail, which can be long and contractile (myophage), long and flexible (siphophage) or short (podophage). The distal extremity of the tail bares the host-recognition apparatus, that also allows the perforation of the bacterium cell wall and the safe channelling of the DNA into the host cytoplasm. In long tailed phages, the assembly pathway of the capsid and of the tail are independent, and DNA-filled capsids attach onto functional tails to form fully assembled virions at the end of the lytic cycle. In siphophages, functional tails are produced after capping of the tail tube by the Tail terminator Proteins (TrP), of which siphophage λ gpU is the prototype (1). In myophages, tails can be capped both by a tube terminator and by a sheath terminator, as in T4 (2), or only by a tail terminator, which caps both the tube and the sheath, as in P1 (3) or Pam3 (4). In some phages, the tail needs the addition of the Tail Completion Protein (TCP) to form fully functional tails, of which λ gpZ is the prototype (5). In some publications, both names are used interchangeably to designate the tail terminator. Empirically, it has been observed that phage proteins within the tail are arranged in a similar order than their coding genes in the genome (5, 6). Thus, it is expected that the TCP, also called Ne1 (Neck protein type 1) in (6), encoded by a gene usually located downstream the capsid module and upstream the TrP of the tail module, itself upstream the Tail Tube Protein, would also be located at the capsid-to-tail interface, the neck (5–7).

Phage T5 that infects *E. coli*, belongs to the morphotype siphophage, which accounts for the majority of tailed phages. In the structure module of its genome, under the control of the same promoter and located between the Head Completion Protein gene and the Tail Tube protein gene, are found two genes, p142 and p143 that were attributed to the TrP and the TCP, respectively (Fig. 1A,H)(8): TrP_p142_, was indeed localised within the virion in the neck using cross-linking with anti-TrP_p142_ IgG and immunogold labelling combined with EM imaging (8). A similar approach for TCP_p143_ proved unsuccessful: TCP_p143_ could be overexpressed in some conditions, but the protein precipitated on the Ni-NTA column, preventing the obtention of purified protein to raise antibodies (9). We recently determined the structure of T5 tail tip – the distal extremity of the tail – before and after interaction with its receptor (10), where we proposed that a monomer of TCP_p143_ be attached to the Baseplate Hub Protein pb3 (BHP_pb3_). Here, we present the cryo-EM structure of phage T5 tail proximal extremity at a pseudo atomic resolution, also before and after interaction with the phage receptor. TrP_p142_ is resolved as a hexameric ring, interacting with the last Tail Tube Protein TTP_pb6_ ring. No available density could account for TCP_p143_ in this region of the tail.

**Figure 1:**
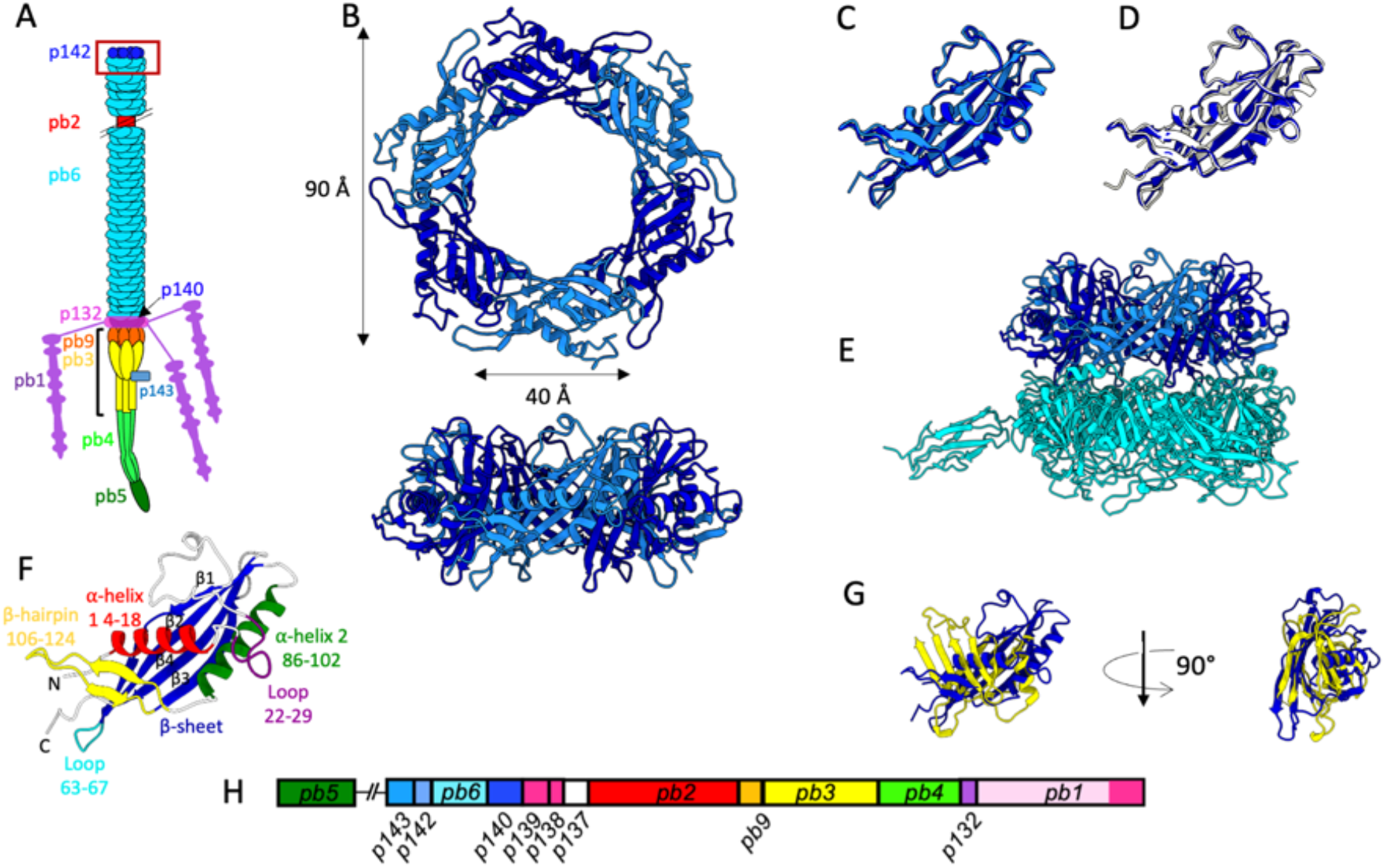
(A) Scheme of T5 tail organisation. The proximal end of the tail (formed of Trp_p142_ and TTP_pb6_ rings) is framed in red and the upper part of the distal complex (comprising TCP_p143_, DTP_pb9_ and BHP_pb3_) is indicated by a black accolade. (B) Top (upper panel) and side (lower panel) views of the Trp_p142_ hexamer as found at the distal extremity of the tail. Two consecutive monomers are not related by a C6 symmetry and are displayed in two different shades of blue. The outer and inner dimensions of the ring are indicated. (C) Superimposition of two Trp_p142_ consecutive monomers (non-symmetry related). The two monomers are nearly identical. (D) Superimposition of two Trp_p142_ monomers from native tails (blue) and tails after interaction with receptor FhuA (white). (E) Side view of the proximal extremity of the tail, featuring the Trp_p142_ hexamer (in shades of blue) sitting on top of the TTP_pb6_ trimer (in turquoise). (F) Annotated model of Trp_p142_, indicating its main secondary structures. The β-sheets are numbered with a black figure above each of them. C and N-termini are also indicated. (G) Superimposition of Trp_p142_ (blue) and a Tail Tube Domain from TTP_pb6_ (yellow). (H) Map of T5 tail structural proteins genes.

## Results

### Structure of the proximal extremity of phage T5 tail

T5 tails, purified from a capsid defective mutant, were produced, vitrified for cryo-EM characterisation and imaged for data collection on a high-end electron microscope (Fig. S1A) (10). Image analysis yielded a map of T5 tail proximal extremity, containing a trimeric ring of the tail TTP_pb6_, on top of which sits a hexameric ring formed by the TrP_p142_ (Fig. 1E). The estimated resolution of this map is 3.9 Å (Fig. S1C,D), which allowed to build pseudo-atomic models for both TrP_p142_ hexamer and a TTP_pb6_ trimer. TrP_p142_ is a 161-residues protein, mainly formed by a 4-strand antiparallel β-sheet that lines the lumen of the tube, flanked by two α-helices, and a short loop between β-strands 1 and 2. An 18-residues β-hairpin links the second α-helix to the β-sheet (Fig. 1F). As already discussed in the case of phage λ TrP gpU (11), TrP_p142_ fold appears to be related to the canonical Tail Tube Domain (TTD) fold, a common building block present in numerous phage proteins (12, 13), although clear differences are present (Fig. S1G). It indeed exhibits a similar β-sheet/helix/loop fold, but a second α-helix is present in TrP_p142_ while a second β-sheet is present in the TTD. Also, the orientation of the helices is different (Fig. 1G). The interaction between two neighbouring TrP_p142_ monomers is mostly mediated through strand β1, N-terminus and C-terminus for monomer 1 and strand β3, α-helix 2 and the 22-29 loop for monomer 2. The interaction between the TrP_p142_ ring and the TTP_pb6_ ring under it is ensured by the tip of the loop 63-67, that inserts in two similar pockets resulting of the domain duplication of TTP_pb6_ (14) and is mostly mediated by electrostatic interactions (Fig. S2). As TrP_p142_ hexameric ring sits on top of a TTP_pb6_ trimeric ring, it exhibits a C3/pseudo-C6 symmetry and two consecutives TrP_p142_ monomers are not strictly identical (Fig. 1B,C). However, as TTP_pb6_ also exhibits a C3/pseudo-C6 symmetry (excluding its Ig-like domain), the two consecutive TrP_p142_ monomers are almost superimposable (rmsd = 1.02 Å over 156 residues). This is reminiscent of what was observed for T5 distal tail protein DTP_pb9_, which also assembles as a hexamer sitting above a trimer (10).

Interestingly, purified TrPs crystallise as biologically relevant hexamers (PDB 3FZB (1) 2GJV and 4ACV). Also, a previous study showed that over-expression of TrP_p142_ with a His_6_-tag in N-terminus lead to soluble, monomeric protein while over-expression of TrP_p142_ with a His_6_-tag in C-terminus (TrP_p142-Cter_) leads to the formation of fibres, of a diameter similar to that of T5 tails (9). This illustrates the fact that phage structural proteins have not only evolved to adopt a specific structure, allowing them to be a building blocks for the phage, but also to bear an inherent dynamic when monomeric. This allows them to remain monomeric, even at high concentrations, and to polymerise and assemble *only* in presence of their partners, to form a fully functional virion. Polymerisation however also occurs when conditions are extreme, as in a crystallisation drop. In the case of TrP_p142-Cter_, the tag addition somehow modifies the dynamics of the protein, which induces aberrant polymerisation upon overexpression. TrP_p142-Cter_ fibres do not look like empty tubes, as would be expected by the piling of TrP_p142_ rings. It could be that the C-terminal tag folds back in the lumen of the tube, filling it and explaining why affinity purification failed (9). Several structures of TTP_pb6_ were previously solved, in different contexts: either as a crystallised monomer, within the tail tube or interacting with the DTP_pb9_ (10, 14). Our study adds the structure of TTP_pb6_ in contact with the TrP_p142_ to these known structures. Overall, there are very little variations between the different TTP_pb6_ structures and TTP_pb6_ in interaction with TrP_p142_ superimposes almost perfectly with TTP_pb6_ interacting with another TTP_pb6_ or with DTP_pb9_ (rmsd of 0.655 Å and 0.576 Å respectively, over the 464 residues).

TrPs form a family of largely conserved proteins amongst sipho- and myophages, in agreement with the fact that DALI searches link Trp_p142_ to several phage TrPs from siphophages as well as from myophages, bacterial tail-like systems related to myo- or siphophages or prophage proteins (Fig. 3A-D, Table S1), with DALI Z-scores ranging from 23.9 to 5.4 and alignment rmsd from 1.2 to 4.2 Å. Several structures of siphophage TrPs are available, either from purified protein or within the phage context: gp17 from SPP1 (15–17), gpU from λ (1, 18– 20)(Fig. 2C,D), gp18 from R4C (21) and gp119 from phage DT57C (22), determined by NMR, X-ray crystallography and/or cryo-EM. *In phago* structures confirm that TrPs indeed assemble as a hexamer, at the proximal extremity of the tail tube, thereby terminating tube polymerisation and providing the surface for interaction with the capsid (17–19, 22). Deletion of this protein has various phenotypes: in phage λ, capsids are produced along with very long, TMP-less, polytails, resulting from the endless polymerisation of the TTP (23); in myophages T4 and P2, deletion mutants of the tube terminators gp3 and gpR, respectively, result in tails with a longer tube, whereas deletion mutants of the sheath terminators gp15 and gpS, respectively, bear longer sheaths (24–26); in myophages Mu and SPO1, deletions mutants of the TrP K and 8, respectively, result in longer tails. In contracted tails, the tubes protrude from the capsid-proximal extremity of the tails rather than from the base plate (27, 28). In siphophage SPP1, normal-length tails are assembled in deletion mutant of the TrP gp17, which however cannot attach to full capsids (29). Thus, TrPs are suggested to both allow to stop the tail polymerisation at the right length, together with the TMP (30), but also to allow interaction with the capsid (29). Some TrPs are larger proteins and include additional domains, for example interacting with the sheath of the myophage tails when they also fill the sheath termination role. Interestingly, PVC16 TrP C-terminus additional domain appears to share structural similarity with the TrP_p142_ fold, indicating domain duplication, a common “LEGO” strategy amongst phages and phage-related machines (13).

**Figure 2:**
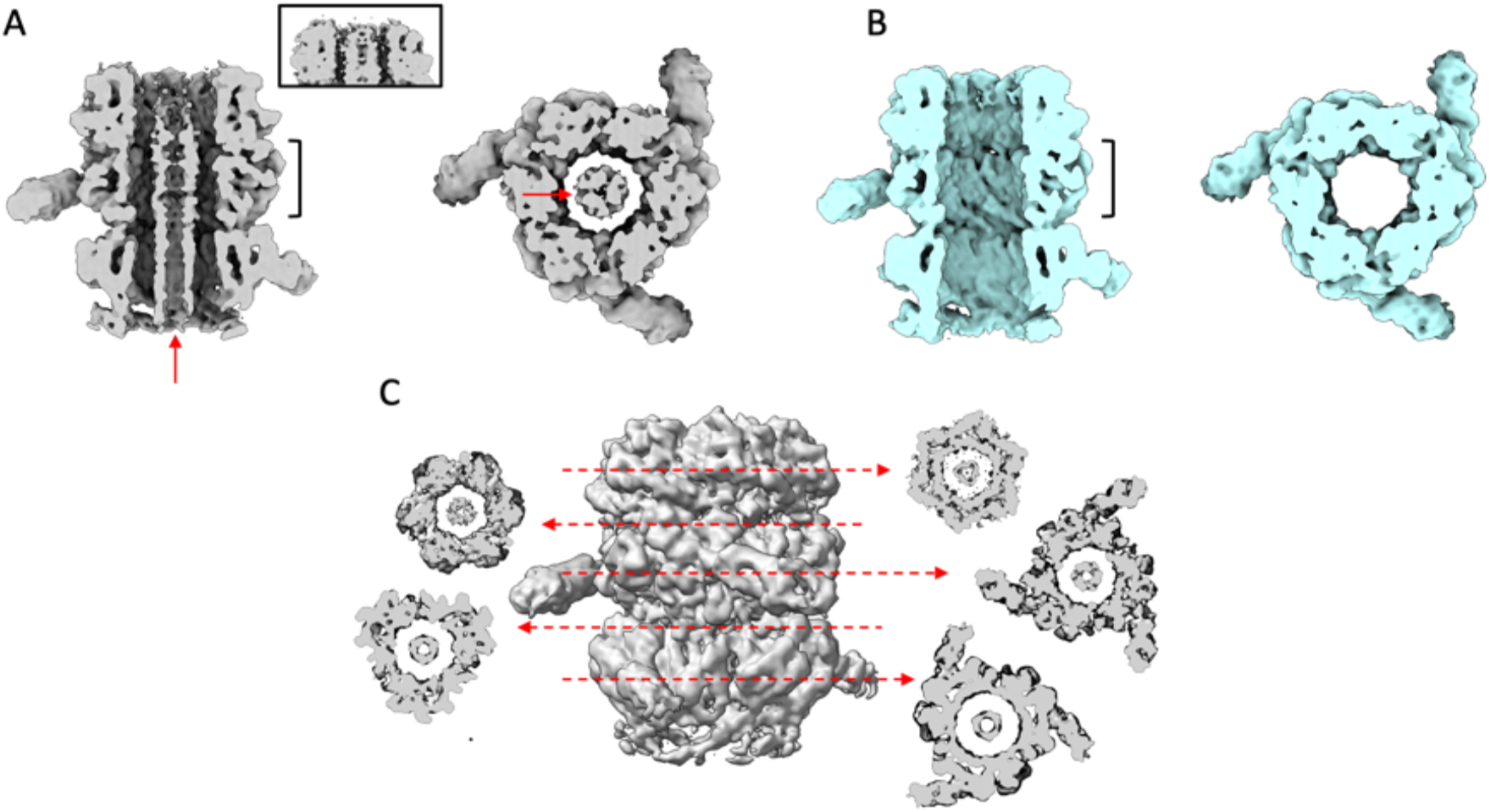
Isosurface view of the non-sharpened cryo-EM maps of T5 tail proximal extremity, in its native state (A, grey) and after interaction with the bacterial receptor FhuA (B, cyan). Left: side interior view and right: top view. The position and thickness of the top view slice is indicated by black accolades in the side view slice. TMP_pb2_ is indicated with red arrows in (A) and is absent in (B). The framed panel in (A) shows the upper part of the tail tube extremity at a higher threshold, with TMP_pb2_ running up to the very end of the tube. (C) Slices of a C3 map of the proximal extremity of T5 tail tube at different positions, showing TMP_pb2_ inside the lumen. All along this section of the tube, TMP_pb2_ appears to be trimeric.

**Figure 3:**
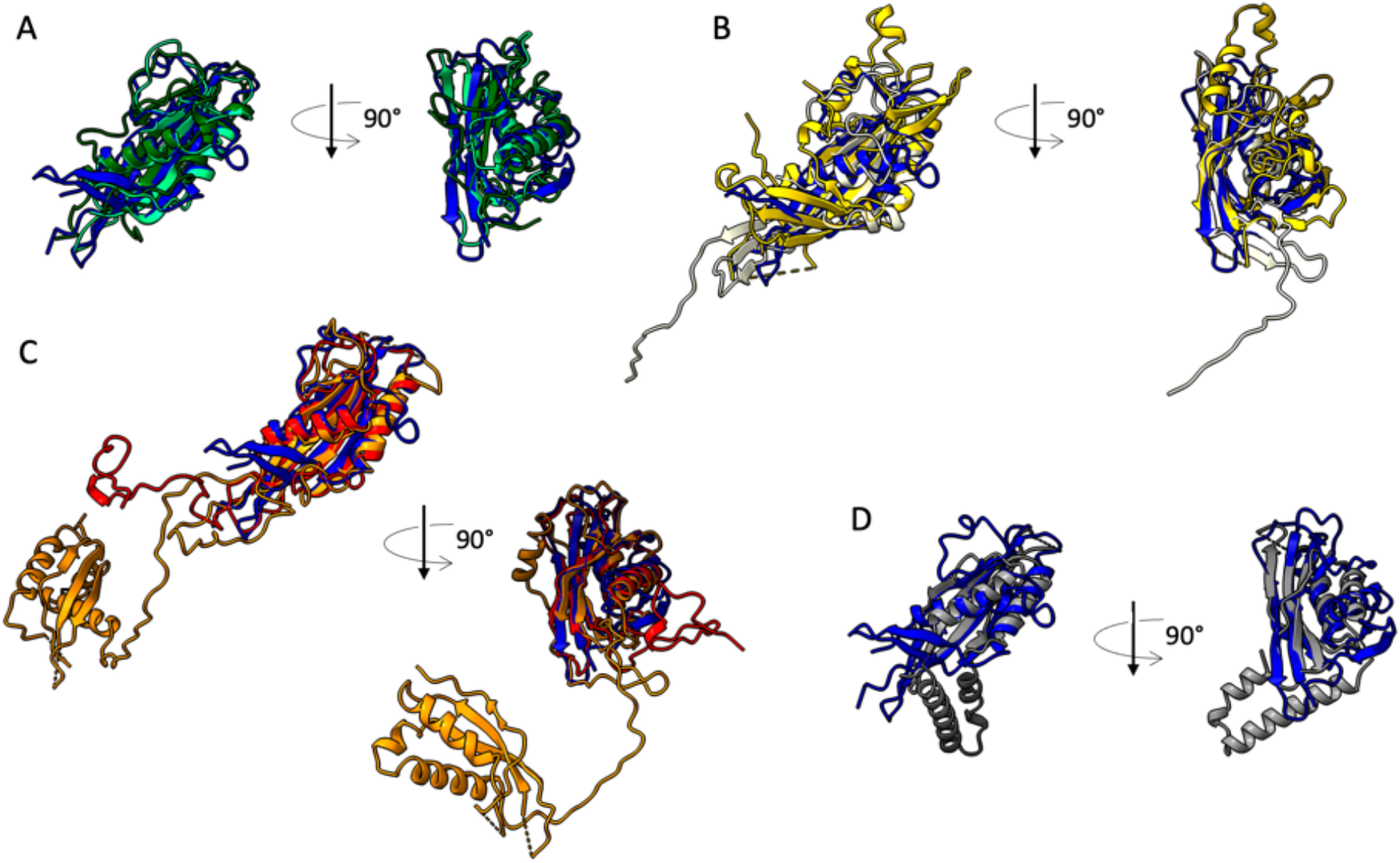
Comparison of TrP_p142_ with TrPs homologous proteins using DALI pairwise. TrP_p142_ is displayed in all panels, coloured in dark blue and superimposed to a selection of homologous proteins of different categories. (A) Siphophages and Siphophage-like machines, phage l (dark green), GTA (light green). (B) Myophages, phage T4 (yellow), phage XM1 (white). (C) Phage-like machines related to Myophages. PyocinR2 (red), PVC (orange). (D) Miscellaneous. Type4 fimbrial biogenesis protein PilO from *P. aeruginosa (*grey).

### The Tape Measure Protein TMP_pb2_

The unsharpened cryo-EM map of T5 tail proximal extremity shows a 25 Å diameter large, cylindrical and ill-defined density filling the tail tube lumen to the top of TrP_p142_, which disappears from the map after interaction with FhuA (Fig. 2A,B). This density most probably belongs to the N-terminus of the Tape Measure Protein pb2 (TMP_pb2_), which fills the entire tail tube in native phages and purified tails (14). The resolution is too low to propose a stoichiometry, yet alone a model, but as a trimer was solved for the C-terminus of the protein (10), is it reasonable to propose that a trimer is also present in N-terminus. Also, and despite the low resolution, different slices taken along the cryo-EM map of T5 tail proximal extremity show a trimeric TMP_pb2_ (Fig. 2C).

TMP stoichiometry has remained uncertain for many decades. In 1973, Zweig and Cummings measured the relative abundance of the different proteins of phage T5, based on a band densitometry analysis of radiolabelled phage proteins after SDS-PAGE (31). TMP_pb2_, BHP_pb3_ and the straight fibre protein pb4 were observed to represent 1.5, 1.5 and 1%, respectively, of the structural proteins of phage T5. In 1988, McCorquodale and Warner interpreted these data and concluded that TMP_pb2_, BHP_pb3_ and pb4 were present as pentamers in the virion (32). In 1974, Casjens and Hendrix proposed a stoichiometry of 6-7 per virion for TMP_gpH_ of phage λ, based on densitometry measurements of phage proteins bands after SDS-PAGE and Coomassie blue or Fast green staining (33). Both in T5 (10, 22) and in λ (18, 19) however, a trimer of the TMP was solved in C-terminus of the protein, in tight interaction with the BHP, as for siphophages 80α (34). For T5, this is consistent with Zweig and Cummings’ results, as BHP_pb3_ and pb4 were also found to be trimers (10). In the case of phage λ, protein staining can quite depend on sequence, making Casjens and Hendrix estimation rather approximative (33). A stoichiometry of six has been however proposed for the TMP for the myophage Pam3 (4). As it is based on a low-resolution map with a C6 axial symmetry applied, it would need to be confirmed.

TMP_pb2_ reaches the proximal extremity of the tube, indicating a possible direct interaction with the DNA in the full phage, as seen in other siphophages (*e*.*g*. SPP1 (15), λ (18)). As previously proposed, and because TMP_pb2_ is expelled upon infection (10, 14), the expulsion of the TMP would be part of the signal transmission pathway that starts with host recognition by the receptor binding protein, continues with cell wall perforation and ends with DNA ejection. All along the tube and except for the 35 last residues in the tail tip, TMP_pb2_ does not interact with the tube wall (10, 14), which can explain the low resolution observed for that protein; this is the case in all phage tail structures determined to date.

### Structure of the proximal extremity of the tail after interaction with receptor FhuA

We also imaged T5 tails after interaction with *E. coli* receptor FhuA reconstituted into nanodiscs (Tail-FhuA) with cryo-EM (10). Image analysis yielded a second 3D map for the proximal extremity of Tail-FhuA, at an average resolution of 4 Å (Fig. S1C,D) allowing to build pseudo-atomic models for TrP_p142_ and TTP_pb6_. At the resolution of our maps, no conformational change was observed in TrP_p142_ monomers or hexamer (rmsd 0.43 Å over 156 residues between Tail and Tail-FhuA individual monomers) (Fig. 1D), and neither in TTP_pb6_. For the latter, it was already shown that T5 tail tube, formed of a stack of TTP_pb6_ trimeric rings, does not undergo any conformational changes upon interaction with FhuA, and that the signal of the infection is not transmitted from the receptor to the capsid through the tube itself (14). TrPs homologous proteins from myophages or tail-like machines also show no conformational change upon infection (2, 35, 36).

Because TMP_pb2_ is expelled during the infection process (8, 10), the lumen of the tube is empty in Tail-FhuA (Fig. 2B), which constitutes the only difference between the two EM maps of the tail proximal extremity. In order to increase the number of particles and to achieve better resolution, we combined both EM datasets but it did not lead to any sensible improvement of the map.

### The tail completion protein TCP_p143_

As mentioned in the introduction, TCPs are a highly conserved family of tail proteins, expected to be located in the neck of phages given the position of their genes in the tail module of phage genomes (5–7). Phage T5 TCP p143, TCP_p143_, is a 27.8 kDa protein, which gene is sandwiched between that of T5 Head Completion Protein and of TrP_p142_ (Fig. 1H). In the cryo-EM map of T5 tail proximal extremity presented here, we could clearly trace TTP_pb6_ trimers and a hexamer of TrP_p142_ (Fig. 1E, 2A-B). Inside the tube, low-resolution density, similar to that present along the whole tube, can reasonably be attributed to TMP_pb2_ (see above). Thus, there is no unassigned density in this proximal extremity map of the tail that could account for TCP_p143_.

TCP_p143_ is however present in the T5 virion and purified tails, as it was unambiguously detected by mass spectrometry (8, 10). We recently determined the high-resolution cryo-EM structure of T5 tail tip. In the C3 map, at the base of BHP_pb3_, a faint density, was visible. Intrigued, we calculated an unsymmetrised map, focusing on this density, and we could indeed solve a *bona fide* density, otherwise blurred by the imposed three-fold symmetry, located at the base of the BHP_pb3_, in the groove between the closed tube and the first fibronectin domain (Fig. 4A, S2A)(10). This density appears only on one of BHP_pb3_ monomers, which is rather unexpected considering the overall C3 symmetry of the rest of the tail. Indeed, the only other monomeric protein of the assembly is the Receptor Binding Protein pb5, located at the extremity of the straight fibre (Fig. 1A)(37). The map resolution in this area (6-10 Å) did not allow to build a protein model *de novo* in this density (Fig. S2A), however, the only unlocalised tail protein remained TCP_p143_.

**Figure 4:**
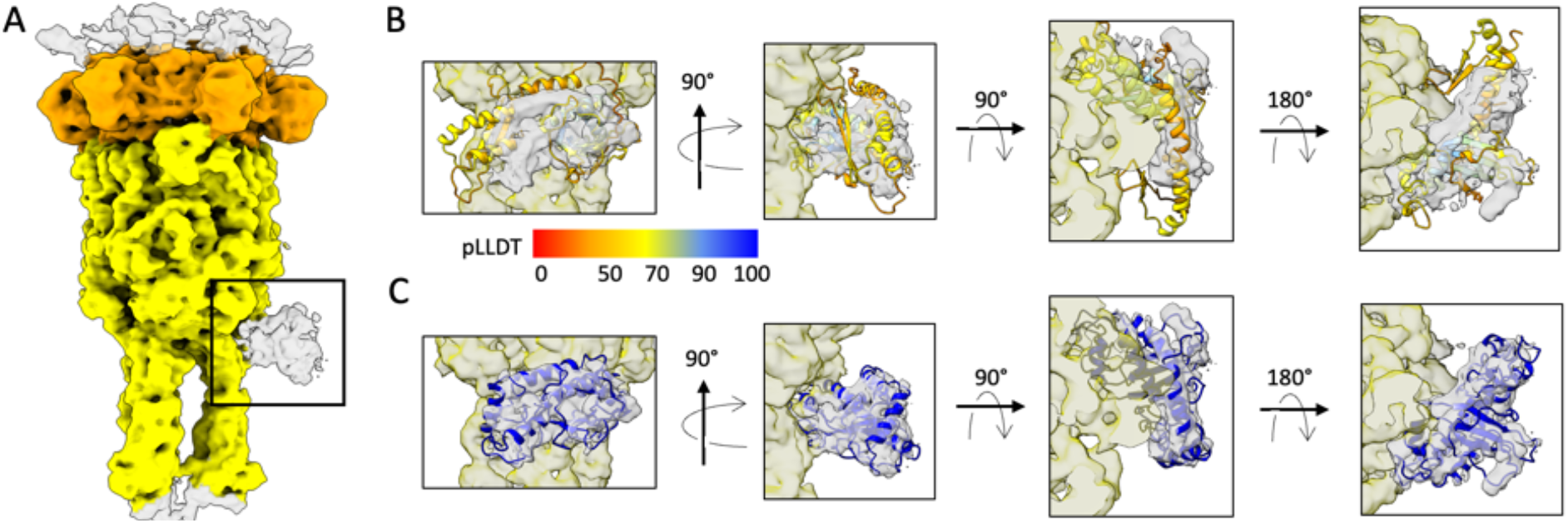
(A) Isosurface view of an unsymmetrised and unsharpened cryo-EM map of the tail tip (side view). The BHP_pb3_ trimer is in yellow and the DTP_pb9_ hexamer is in orange. The additional density located on the side of the BHP_pb3_ trimer and corresponding to the monomeric protein TCP_p143_ is framed in black. (B-C) Fit of an AF2 predicted model in the additional densities ((B) unmodified AF2 model, (C) flexible-fitted AF2 model, coloured blue). Different views (from left to right: front view, side view, top slice, bottom slice) are shown. The model in panel (B) is coloured according to its per-residue pLLDT / confidence measure. Colour key interpretation is the following: 100 to 90 = high accuracy expected, 90 to 70 = backbone expected to be correctly modelled, 70 to 50 = low confidence/caution, 50 to 0 = should not be interpreted, strong predictor of disordered regions.

We used Alphafold2 (AF2) (38) to predict TCP_p143_ structure and see whether it could be fitted into the unassigned density. The five AF2 predicted TCP_p143_ structures are rather heterogeneous and exhibit relatively low confidence levels (pLLDT between 30 and 90%, Fig. 5, S3A), which could indicate a highly flexible protein. AF2-TCP_p143_ seems however to be composed of two domains (residues 1-80 and 81-255)(Fig. S3B-C). The global size and shape of some of the AF2 predicted models correspond to the size and shape of the unattributed density (Fig. 4B) and flexible fitting led to a convincing fit of the predicted secondary structures (Fig. 4C, Suppl. file). The fitted structure is mostly composed of α-helices and loops, with a small β-sheet formed by 3 short strands (Fig. 5, S3A). Thus, we propose that this unallocated density indeed belongs to a monomer of TCP_p143_. Even though our results indicate that only one TCP_p143_ copy per phage is present, cryo-EM single particle analysis consisting in the averaging of a large number of individual particles, we cannot exclude the possibilities of zero, two or three TCP_p143_ copies being present on T5 tails. If it were the case though, these occurrences would only account for a minority of particles, as there is no hint of more than one full occupancy in our unsymmetrised map. We cannot explain yet why only one TCP_p143_ molecule would bind to T5 tail tip, as BHP_pb3_ trimer follows a C3 symmetry and that, at the resolution of our map, no difference in the three potential binding sites could be identified (Fig. S2B-D).

**Figure 5:**
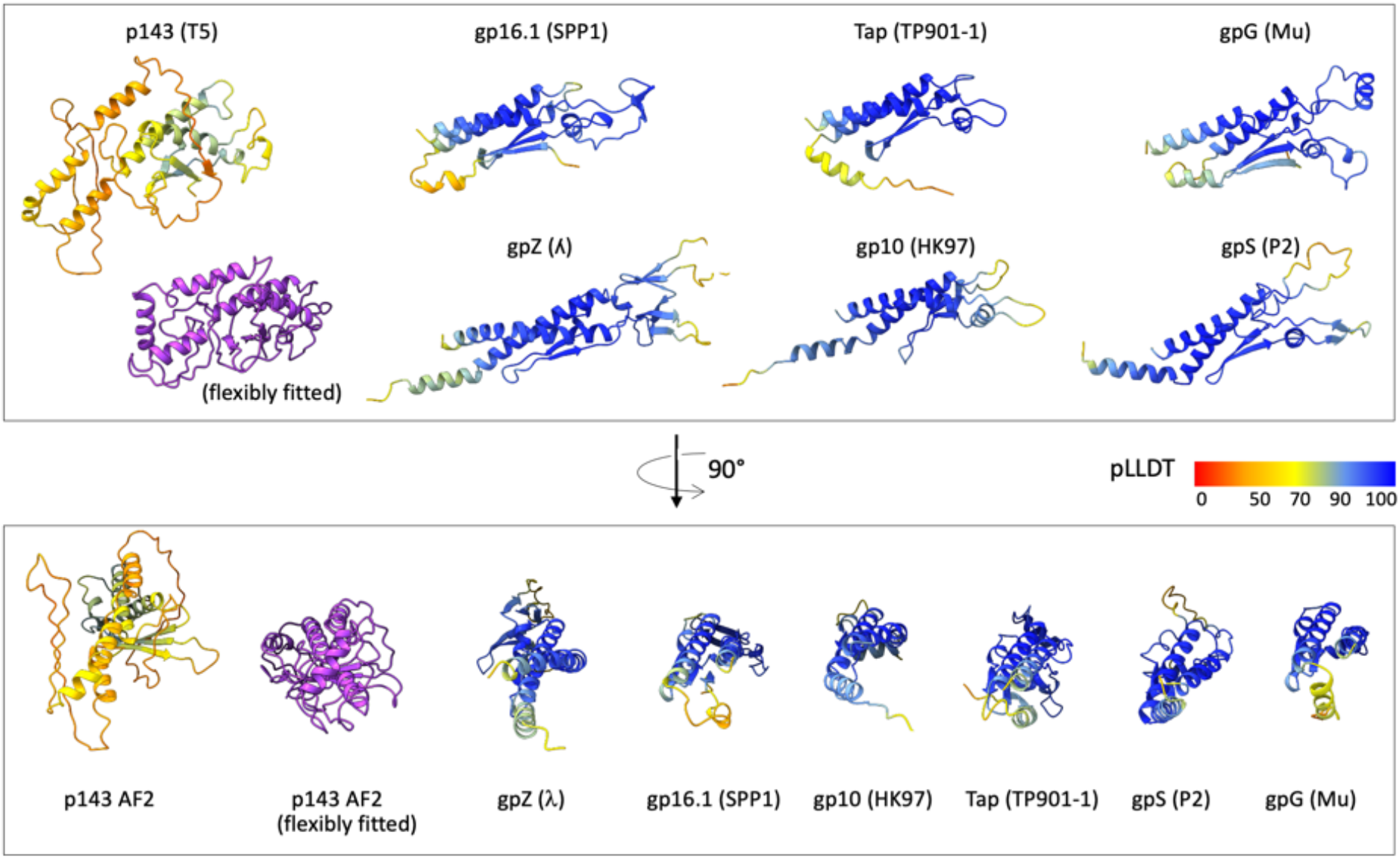
**Alphafold2 predictions for different phage TCPs**, coloured according to their per-residue pLLDT (confidence measure), except for the flexible fitted TCP_p143_ (purple). The pLLDT colour key interpretation is as in Figure 3. The bottom panel models are rotated horizontally by 90° compared to the ones of the top panel. Uniprot entries: T5 (Q6QGE0), λ (P03731), SPP1 (O48447), HK97 (E9RJ97), TP901-1 (Q77K21), P2 (P36934) and Mu (Q01261).

In the EM map of T5 tail tip after interaction with its receptor FhuA, TCP_p143_ is absent (Fig. S2E). This is not surprising given the position of TCP_p143_ at the interface between the tip of the tube and the straight fibre, as the large conformational changes induced by receptor binding, inducing the bending of the straight fibre on the side of the tube, completely re-arrange/destroy TCP_p143_ binding site (Fig. S2E). Given the asymmetry induced by TCP_p143_ in the tail tip, its presence could determine the direction in which the straight fibre bends. In any case, it would definitely detach upon infection and could play an active role, remaining to be determined, in the infection process.

To see whether the putative TCPs of different phages share more than a common position in the structural gene module in phage genomes, we predicted the structures of some TCPs (Fig. 5). A first striking difference with TCP_p143_ is that all structures are predicted with a high level of confidence (pLLDT > 90 for most of the protein). A second interesting feature is that, even though varying in length, from 112 residues for TP901-1 TCP to 192 for λ TCP, these predicted structures all share common features, including two side-by-side long α-helices and a small β-sheet, and superimpose very well (Fig. S3D). AF2-TCP_p143_ seems to stand out from the other predicted TCPs in terms of structure and length (255 residues), and thus, it could maybe also stand out in terms of location and be located at a different position in the tail.

As far as we know, no structure of a phage TCP has been solved to date, and their function and localisation within the tail remain unknown. Literature about TCPs is scarce, contradictory and does not help to set the case about their localisation or function. Indeed, TCP deletion sometimes leads to phages failing to assemble correctly (tail and capsid are produced but do not assemble) as in myophage P2 (26), siphophages TP901-1 (39) and SPP1 (40). In other phages, like siphophage λ (gpZ) (33) or myophage Mu (27), TCP deletion leads to normal-looking virions that however exhibit much lower infectivity. Such a dramatic functional difference is difficult to explain, especially considering that their predicted structures appear very similar. The different phenotype of TCP deleted mutants however does not stem from a different neck organisation between siphophages and myophages, as in TCP less mutants, myophages Mu and P2 behave differently, as well as siphophages SPP1, TP-901-1 and λ. In the case of phages P2, TP901-1 and SPP1, the fact that TCP deleted mutants do not form complete virions suggests that the presence of this protein is a prerequisite to capsid attachment to the tail, and the most straightforward conclusion would be that it is located in the neck region (40). However, the structure of SPP1 neck has been determined and, as for T5, unambiguously shows that its TCP-gp16.1 is not present (17). This is also the case for λ, in which TCP-gpZ is detected in the tail particle but not localised in the tail structure (18, 19), and for all myophage neck structures solved to date, where there is no trace of a protein that would be a TCP (*e*.*g*. 4, 40, 41). In λ, in which full virions are produced in a TCP-deleted mutant, it was suggested that TCP-gpZ allows for the correct DNA insertion into the tail after capsid and tail attachment (43), explaining the lack of infectivity of gpZ-deleted virions. TCP_p143_ is a small, monomeric protein, part of a highly symmetrical megadalton-sized complex, with a very unexpected location. Thus, special attention and sophisticated image processing was required to reveal it. We would like to kindly encourage our colleagues to have a fresh look at their phage tail cryo-EM data, as this protein should also be present in a majority of phage tails. Increasing data would allow to shed light on this still mysterious protein.

## Conclusion

We present the structure of the proximal extremity of phage T5 tail. In the densities available, we could trace, as expected, a hexamer of TrP_p142_ and a first trimer of TTP_pb6_. In the lumen of the tail tube before interaction with the receptor, densities of much lower resolution are present, which we attribute to TMP_pb2_. As for the many structures of proximal tail extremity or necks of either sipho- or myophages, no remaining densities, even faint, could correspond to the TCP. Instead, we could localise a medium-resolution unattributed density in the unsymmetrised map of the distal extremity of T5 tail, at the base of a BHP_pb3_ monomer, in which a predicted structure of TCP_p143_ could be fitted. Predicted structures of other sipho- and myophages TCPs share the same fold, whereas TCP_p143_ seem to stand out in several aspects. Given the dispersity of phenotypes of TCP deleted phages, it is thus possible that this protein adopts different roles and location within different phage tails.

## Acknowledgments

We thank Guy Schoehn for the general scientific and instrumentation support, Grégory Effantin for cryo-EM data collection, Emmanuelle Neumann, Daphna Fenel and Séraphine Degroux for technical assistance, Leandro Estrozi, Ambroise Desfosses, Benoît Arragain for help and discussion on image analysis and Aymeric Peuch for help with the use of the EM computing cluster. We acknowledge the European Synchrotron Radiation Facility for provision of beam time on CM01. This work used the platforms of the Grenoble Instruct-ERIC center (ISBG; UMS 3518 CNRS-CEA-UGA-EMBL) within the Grenoble Partnership for Structural Biology (PSB), supported by FRISBI (ANR-10-INBS-05-02) and GRAL, financed within the University Grenoble Alpes graduate school (Ecoles Universitaires de Recherche) CBH-EUR-GS (ANR-17-EURE-0003). The IBS acknowledges integration into the Interdisciplinary Research Institute of Grenoble (IRIG, CEA).

## Author Contributions

CB conceived the project. RL and CB prepared the T5 tails and FhuA-nanodiscs used for high-resolution cryo-EM. RL optimised cryo-grid preparation, processed the cryo-EM data, built the atomic models, analysed and interpreted the models with the help of CB. RL and CB wrote the paper.

## Funding

This research was funded by the Agence Nationale de la Recherche, grant number Projet ANR-21-CE11-0023. The electron microscope facility is supported by the Auvergne-Rhône-Alpes Region, the Fondation pour la Recherche Médicale (FRM), the Fonds FEDER and the GIS-Infrastructures en Biologie Santé et Agronomie (IBiSA).

## Conflicts of Interest

The authors declare no conflict of interest.

## Data Availability

The cryo-EM density maps of T5 tail proximal extremity presented in this study and the associated atomic coordinates have been respectively deposited in the Electron Microscopy Data Bank (EMDB) and in the Protein Data Bank (PDB) under the following accession codes: EMD-15967 / PDB 8BCP (native state) and EMD-15968 / PDB 8BCU (after interaction with FhuA). The unsymmetrised cryo-EM density map of T5 tail tip has been deposited in the EMDB under accession code EMD-51232. TCP_p143_ AF2 flexible-fitted model is provided as supplementary material.

## Material and Methods

### T5 tail purification

Tails were purified as in (8, 10). Briefly, an *E. coli* strain F culture was infected with the mutant phage T5D20*am*30d, which bears an Amber mutation in the major capsid protein gene. After complete cell lysis and DNA digestion by the addition of DNase, cell debris and unlysed cells were removed by low-speed centrifugation and T5 tails were precipitated by incubation with 0.5 M NaCl and 10 % (w/w) PEG 6000 overnight at 4°C. After low-speed centrifugation, the pellet was resolubilised in buffer and purified on a glycerol step gradient. The gradient fractions containing the tails were diluted ten times, loaded onto an ion exchange column, and the tails were eluted by a 0 – 0.5 M NaCl linear gradient. Purified tails were incubated 30 min with FhuA loaded nanodiscs in a tail:FhuA-nanodisc ratio of 1:10 (vol/vol) at room temperature (10).

### Cryo-EM sample preparation

Typically, 3.5 μL of T5 tails sample (with or without FhuA-nanodisc) were deposited on a freshly glow discharged (25 mA, 30 sec) Cu/Rh 300 mesh Quantifoil R 2/1 EM grids and plunge-frozen in nitrogen-cooled liquid ethane using a ThermoFisher Mark IV Vitrobot device (100% humidity, 20°C, 5 s blotting time, blot force 0).

### EM data acquisition

Respectively 3208 and 5752 micrographs were collected for the tails alone and the FhuA-nanodisc incubated tails, over three different data collections. 40-frame movies were acquired on a ThermoFisher Scientific Titan Krios G3 TEM (European Synchrotron Radiation Facility, Grenoble, France) (44) operated at 300 kV and equipped with a Gatan Quantum energy filter coupled to a Gatan K2 summit direct electron detector. Automated data collection was performed using Thermo Fisher Scientific EPU software, with a typical defocus range of -1.0 to -3.0 μm and a total dose of 40 e^-^/Å^2^ per movie. A nominal magnification of 105.000x was used, resulting in a calibrated pixel size at the specimen level of 1.351 Å. The images used were the same as those used in (10) (Table S2).

### EM image processing

#### Preprocessing and particle picking

Frame alignment was performed using Motioncor2 (45) keeping respectively frames 3 to 30 and 1 to 40 for Tail and Tail, and applying dose weighting. Contrast Transfer Function parameters were then determined using Gctf (46); Manual particle picking was performed with EMAN2/e2helixboxer (47). The picking coordinates were consistently centred on the last visible ring at the proximal extremity of the tail tube, corresponding to the tail terminator p142 hexamer. All subsequent image processing was performed using Relion (versions 3.0 and 3.1) (48) (Table S2).

#### Proximal extremity of the tail (Tail and Tail-FhuA)

Tail and Tail-FhuA data was processed in an identical fashion. After particle extraction (box size of 140×140 pixels^2^), a run of 2D classification was performed to get rid of bad particles (Fig. S1B). A run of 3D classification was then performed, using a cylinder (diameter 90 Å, height 188 Å) as an initial model, at the end of which a small additional fraction of the particles was discarded, ending up with homogeneous dataset of respectively 9,953 and 10,701 particles. 3D refinement with masking and C3 symmetry imposed followed, using the best 3D class as an initial model (low-pass filtered at 15 Å) yielded a C3 map of the proximal extremity of the tail, comprising the tail terminator p142 hexamer and the first TTP_pb6_ ring. After masking and sharpening, the overall estimated resolution of the maps respectively reached 3.88 Å and 4.05 Å (FSC_0.143_) for Tail and Tail-FhuA (Fig. S1D). For both maps, a local resolution map was calculated using Relion built-in local resolution tool (Fig. S1C).

#### Tail completion protein TCP_p143_

The full protocol used to obtain the cryo-EM maps for T5 tip complex is detailed in Linares *et al*. (10). Here, we detail the steps that are specific to the map shown in this study. After particle extraction under T5 collar and 2D classification, a homogeneous dataset of 9290 particles was obtained. No 3D classification was performed. Using a 15-Å resolution map determined from a previous cryo-EM data collection (unpublished) as an initial model, a C3 reconstruction of the tip was calculated, from the second TTP_pb6_ ring to the beginning of the central fibre. Refined particles from the previous reconstruction were reextracted (box size of 200 pixels by 200 pixels) and recentered on the lower part of BHP_pb3_, on the side of which TCP_p143_ is located. After reclassification/selection, symmetry relaxation, and a new run of 3D refinement using suitable masking and no symmetry imposed, a new map of the central part of the tip complex was obtained, where densities for the monomeric TCP_p143_ were visible.

#### Protein model building

Atomic protein models were built into the two cryo-EM maps using the Coot software (49) by tracing the protein sequence into the densities and were then iteratively refined alternating Coot manual refinement and PHENIX (50) real space refine tool until convergence. For Trp_p142_, the model of the homologous Trp-gpU from siphophage λ (PDB: F3Z2) (1) was rigid-body fitted in the cryo-EM map of the proximal extremity of the tail and was used as a starting point for model building, while for TTP_pb6_ models, existing cryo-EM models (PDB: 7QG9) (10) were used as a starting point and were refined into the EM maps. Molprobity (51) was used for model quality assessment (Table S2).

All the predicted structures in this study were generated using AlphaFold2 (38) on ColabFold (https://colab.research.google.com/github/sokrypton/ColabFold/blob/main/AlphaFold2.ipynb) (52). In addition, for TCP_p143_, we fitted the most promising AF2 predicted model into the unsymmetrised tip cryo-EM map, then used a combination of Flex-EM (53) / Namdinator (54) (flexible fitting) and PHENIX (50) (real space refine), in an iterative way, to obtain a better model for this protein, with a convincing fit of most of its secondary structures.

## Supplementary tables and figures

**Table S1:** Table of the relevant DALI hits for T5 Trp_p142_.

| PDB code | Z-score | rmsd | lali | nres | %id | Protein name | Structure determination method |
| --- | --- | --- | --- | --- | --- | --- | --- |
| 8HQO | 23.9 | 1.2 | 158 | 160 | 97 | gp119 from siphophage DT57A* | Cryo-EM, on the full phage |
| 6TOA | 14 | 3 | 128 | 134 | 8 | Adaptor protein RCC01688 from GTA (empty) | Cryo-EM, on the full particle |
| 6TE9 | 14 | 3 | 131 | 134 | 8 | Adaptor protein RCC01688 from GTA (native) | Cryo-EM, on the full particle |
| 2GJV | 13 | 3 | 128 | 136 | 9 | Putative cytoplasmic protein STM4215 (Salmonella prosiphophage) | X-ray diffraction, crystallised as a hexamer |
| 4ACV | 13 | 2 | 115 | 119 | 7 | "Antigen B", prophage $\lambda$ LM01 | X-ray diffraction, crystallised as a hexamer |
| 6U5G | 13 | 3 | 130 | 164 | 8 | Collar protein PA0615 from Pyocin R2 | Cryo-EM, on the full particle |
| 3FZ2 | 13 | 3 | 128 | 131 | 16 | TrP gpU from siphophage $\lambda$ | X-ray diffraction, crystallised as a hexamer |
| 8K37 | 12.3 | 2.7 | 125 | 131 | 18 | TrP gpU from siphophage $\lambda$ | Cryo-EM, on the full phage |
| 7KJK | 10 | 3 | 120 | 159 | 10 | Tail terminator gp5 from XM1 | Cryo-EM, on the full phage |
| 6JOF | 10 | 4 | 127 | 272 | 9 | Pvc16 cap protein from PVC (extended state) | Cryo-EM, on the full particle |
| 3J2N | 9,7 | 4 | 131 | 211 | 11 | Tail connector protein gp15 from myophage T4 (contracted state) | Cryo-EM (25 Å resolution) + X-ray diffraction structure (PDB: 4HUD) |
| 3J2M | 9,3 | 4 | 136 | 211 | 11 | Tail connector protein gp15 from myophage T4 (extended state) | Cryo-EM (25 Å resolution) + X-ray diffraction structure (PDB: 4HUD) |
| 4HUD | 9,4 | 4 | 131 | 211 | 11 | Tail connector protein gp15 from myophage T4 | X-ray diffraction, crystallised as a hexamer |
| 2L25 | 9,2 | 4 | 129 | 141 | 12 | Bordetella Bronchisepta Phage Related Protein | NMR, soluble monomer |
| 6RAP | 9 | 3 | 122 | 275 | 8 | AFP1 protein from AFP *Serratia entomophila* | Cryo-EM, on the full particle |
| 2LFP | 8,7 | 3 | 114 | 139 | 11 | Tail-to-head joining protein gp17 from siphophage SPP1 | Cryo-EM (7-8 Å resolution, PDB 5A20 & 5A21) + NMR structure of the soluble monomer |
| 1Z1Z | 7,6 | 3 | 108 | 129 | 11 | TrP protein gpU from siphophage $\lambda$ | NMR, soluble monomer |
| 5N7L | 5,4 | 3 | 76 | 80 | 11 | Protein L (GspL) from Type II Secretion System | X-ray diffraction |
| 2RJZ | 6,3 | 3 | 85 | 130 | 7 | Type 4 fimbrial biogenesis protein, *P. aeruginosa* | X-ray diffraction |
| 8HDR |  |  |  |  |  | Trp gp12 from myophage Pam3 | Cryo-EM, on the full phage |
| 8GTF |  |  |  |  |  | Trp vBDshSR4C_010 from myophage Dinoroseobacter vB_DshS-R4C | Cryo-EM, on the full phage |
| 8FVH |  |  |  |  |  | Gateway protein gp29 from Pseudomonas phage E217 | Cryo-EM, on the full phage |

\*Protein gp119 from DT57C in (22) is called TCP and is wrongly indicated as TCP ORF T5.147 homologue. It is homologous to TrP ORF T5.146, which codes for TrP<sub>p142</sub>.

**Table S2:** Validation statistics and model building.

|  | Proximal extremity of T5<br>tail<br>(EMD-15967)(PDB 8BCP) | Proximal extremity of T5 tail<br>after interaction with FhuA<br>(EMD-15968) (PDB 8BCU) |
| --- | --- | --- |
| **Data collection and processing** |  |  |
| Magnification | 105.000 x | 105.000 x |
| Voltage (kV) | 300 | 300 |
| Electron exposure (e<sup>-</sup>/Å<sup>2</sup>) | 40 | 40 |
| Defocus range (μm) | -1.0 to -3.0 | -1.0 to -3.0 |
| Pixel size (Å) | 1.351 | 1.351 |
| Symmetry imposed | C3 | C3 |
| Micrographs (no.) | 3208 | 5752 |
| Final particle images (no.) | 9953 | 10701 |
| Map resolution (Å) 0.143 | 3.88 | 4.05 |
| FSC threshold |  |  |
| Map resolution range (Å) | 3.7 – 7 | 4 – 7 |
| **Refinement** |  |  |
| Model resolution (Å) 0.5 | 4.1 | 4.1 |
| FSC threshold |  |  |
| Map sharpening *B* factor (Å<sup>2</sup>) | -146 | -120 |
| **Model composition** |  |  |
| Chain count | 9 | 9 |
| Non-hydrogen atoms | 18171 | 18279 |
| Protein residues | 2337 | 2349 |
| Ligands | 0 | 0 |
| ***B* factors (Å<sup>2</sup>)** |  |  |
| Protein (min/max/mean) | 43.48/297.70/89.1 | 65.51/243.36/101.07 |
| **R.m.s. deviations** |  |  |
| Bond lengths (Å) | 0.006 | 0.005 |
| Bond angles (°) | 1.005 | 0.971 |
| **Validation** |  |  |
| MolProbity score | 1.97 | 2.04 |
| Clashscore | 10.41 | 12.32 |
| Poor rotamers (%) | 0 | 0.15 |
| **Ramachandran plot** |  |  |
| Favoured (%) | 96.51 | 93.26 |
| Allowed (%) | 3.49 | 6.61 |
| Disallowed (%) | 0 | 0.13 |

**Figure S1:**
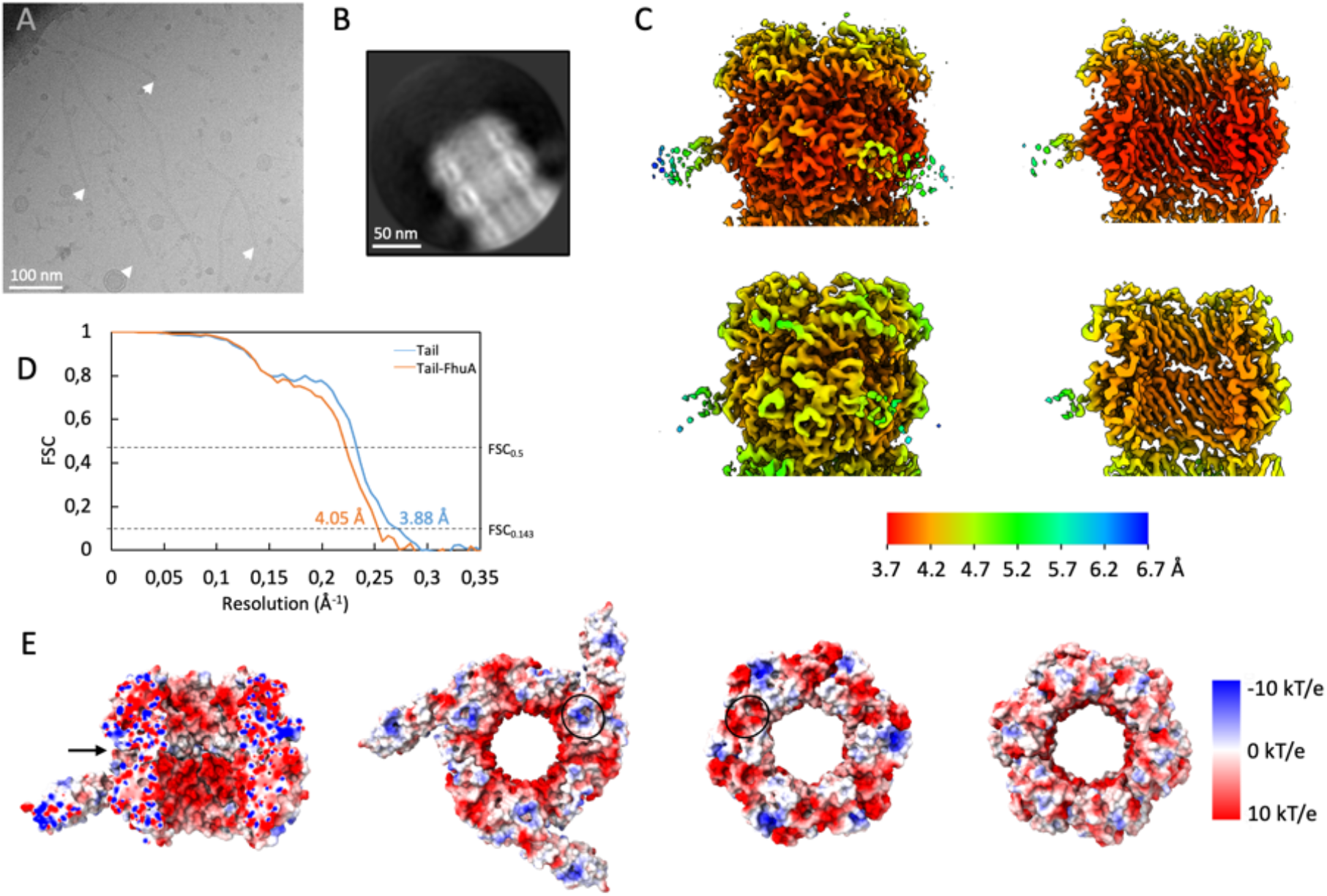
(A) Cryo-EM micrograph of T5 tails. The Trp_p142_ hexamer forms the last ring of the proximal extremity of T5 tail and is indicated with a white arrow. (B) 2D class of T5 tail proximal extremity, side view. (C) Local resolution of the cryo-EM maps for T5 tail proximal extremities, as determined by Relion, in the tail native state (top) and after interaction with *E. coli* receptor FhuA (bottom). The left panel shows a side view for the two maps while the right panel presents an interior side view. All maps were calculated with a C3 symmetry imposed and the key is the same for both maps. Local resolution is the highest at the centre, falls off radially and is the worst at the tip of the flexible TTP_pb6_ Ig-like domain. The top of the Trp_p142_ hexamer appears a bit less resolved, probably because of some flexibility of the binding interface to the Head Completion Protein of the capsid (absent here). (D) Fourier shell correlation (FSC) plot for the maps presented in (C). FSC_0.5_ and FSC_0.143_ cutoffs are indicated, as well as the estimated resolution (FSC_0.143_) for each map. (E) Electrostatic charge distribution of the proximal extremity of T5 tail tube. From left to right: slice from a side view showing the tube lumen (panel 1), top view of the last TTP_pb6_ ring (interacting with TrP_p142_ ring, panel 2), bottom view of TrP_p142_ ring (interacting with TTP_pb6_, panel 3) and top view of the p142 ring (directed toward the capsid, panel 4). Complementary charge patches are circled in black. The position of the TTP_pb6_ –TrP_p142_ interface is indicated with a black arrow in the first panel. Electrostatic charge distribution was calculated with the “coulombic” tool from ChimeraX. Values of the electrostatic charges range from -10 kT/e (blue) to 10 kT/e (red).

**Figure S2:**
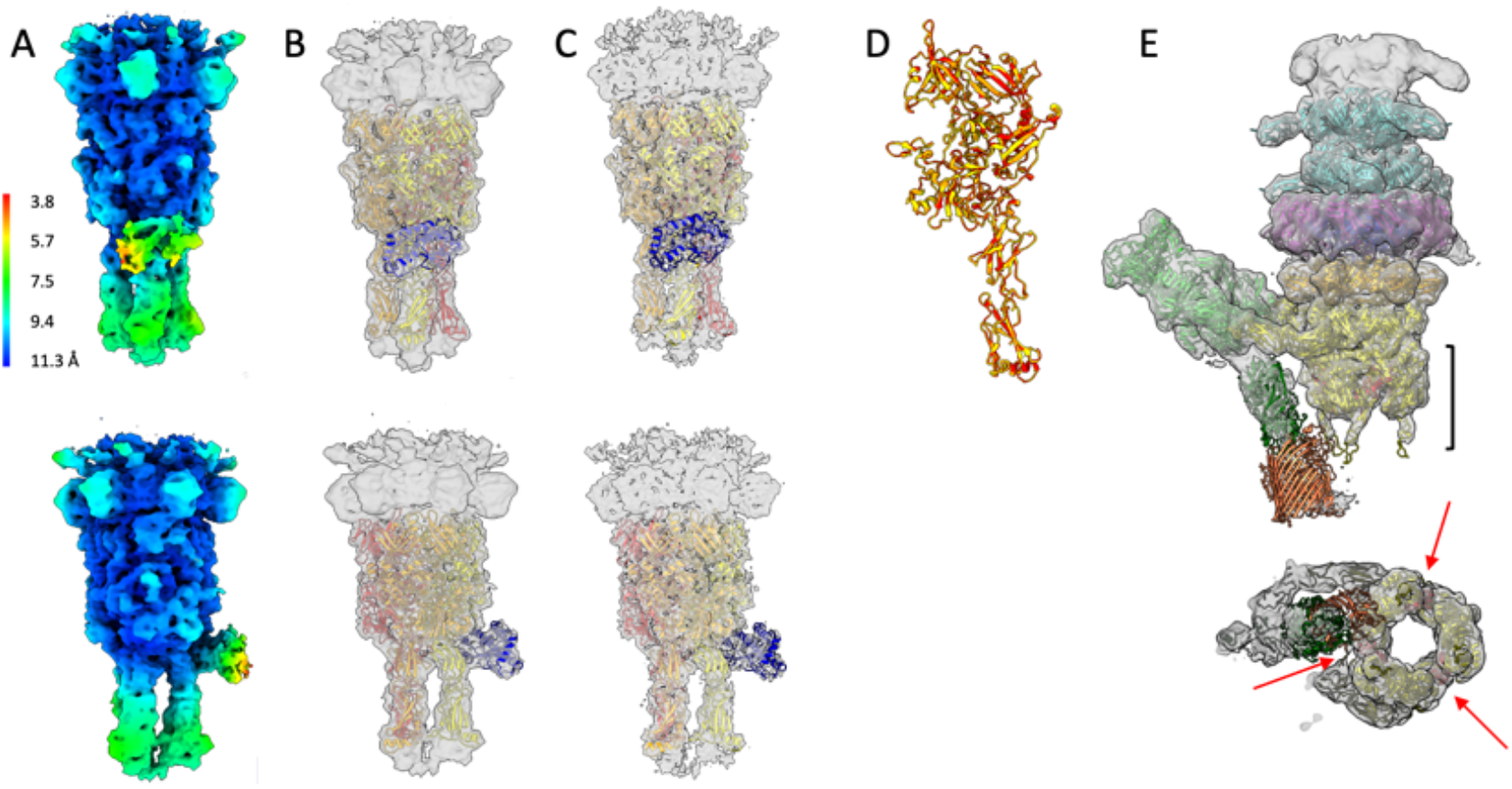
(A) Local resolution of the unsymmetrised cryo-EM map for T5 tip, including the monomeric TCP_p143_, as determined by Relion, front (top) and side (bottom) views. (B-C) Unsharpened (B) and sharpened (C) unsymmetrised EM maps of T5 tip, fitted with three monomers of BHP_pb3_ (yellow, orange, red) and the monomeric TCP_p143_ (blue). (D) Superimposition of the three BHP_pb3_ monomers, individually refined in the unsymmetrised sharpened map of T5 tip shown in (C). rmsd between monomer 1 and 2 or 1 and 3 is respectively 0.368 Å and 0.353 Å over 949 residues. (E) EM map and models of T5 tail tip complex after its interaction with the bacterial receptor FhuA inserted in nanodisc (described in (10)). TCP_p143_ is absent and all the densities are attributed to other proteins, indicating TCP_p143_ would probably detach during the infection process. The location and thickness of the bottom slice is indicated on the first panel by a black accolade. Arrows are pointing to TCP_p143_ position in the map before interaction with the receptor.

**Figure S3:**
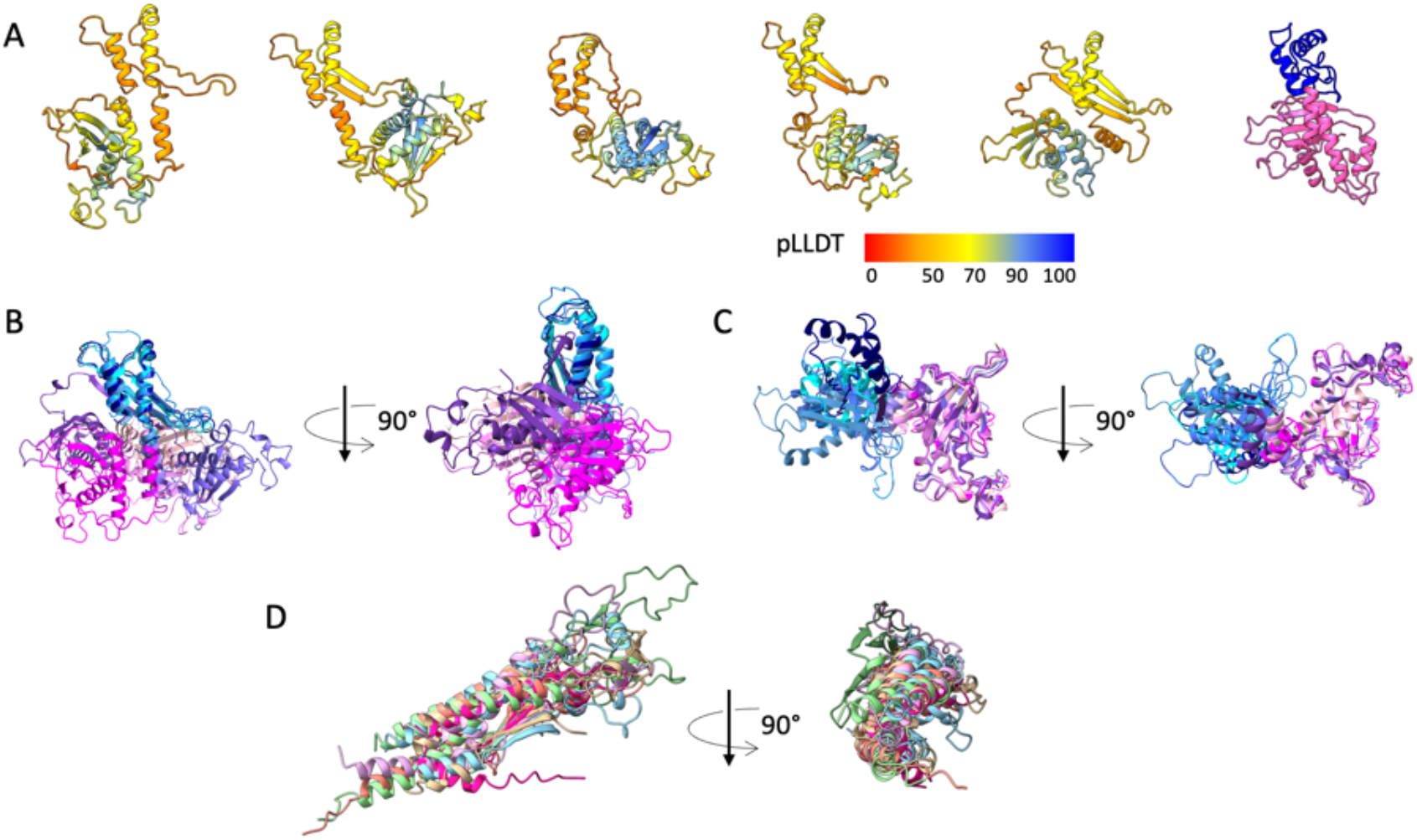
(A) Five different AF2 predictions for T5 TCP_p143_, coloured according to their per-residue pLLDT (confidence measure). The sixth model of the panel is AF2 TCP_p143_ predicted model after flexible fitting in the T5 tip map, with residues 1-80 and 81-255 respectively coloured in blue and pink. (B, C) Superimposition of the five TCP_p143_ models shown in (A), after alignment on residues 1-80 (B, rmsd between 65 pruned atom pairs is 1.057 Å, across all 80 pairs: 1.958) or residues 81-255 (C, rmsd between 137 pruned atom pairs is 0.669 Å, across all 175 pairs: 3.996) using the Matchmaker tool of ChimeraX. Residues 1-80 are coloured in shades of blue and residues 81-255 in shades of pink. (D) DALI pairwise alignment of AF2 predictions for TCPs from phages λ (green), SPP1 (beige), HK97 (orange), TP901-1 (purple), P2 (pink) and Mu (blue).

## Notes

### Competing Interest Statement

The authors have declared no competing interest.

